# Distinct neural modes carry information about grasp force and phase in the sensorimotor cortex

**DOI:** 10.64898/2026.02.01.702680

**Authors:** G.H. Blumenthal, B.M. Dekleva, C. Gontier, I.C. Gonzalez, J.A. Gonzalez-Martinez, B.M. Yu, A.P. Batista, A.R. Sobinov, L.E. Miller, R.A. Gaunt, M.L. Boninger, S.M. Chase, J.L. Collinger

## Abstract

Humans perform a variety of complex hand movements to manipulate objects, requiring precise control of changing forces. Understanding the role of sensorimotor cortex and the cortical dynamics underlying these actions is crucial for developing interventions that restore dexterous hand function after injury or disease. In this study, two individuals with tetraplegia resulting from cervical spinal cord injury attempted a series of isometric grasps. Neural activity was recorded from the motor and somatosensory cortices using intracortical microelectrode arrays while participants attempted to exert a static force or to ramp force up and down. Despite their inability to execute movement, and with limited afferent input, the spiking activity in motor and somatosensory cortex was modulated with the task. Within the neural response we identified independent neural modes - distinct patterns of population-level neural activity - that were informative about both the timing and magnitude of the force. Moreover, distinct neural modes were observed during static and dynamic grasping conditions, suggesting independent control schemes for maintaining and changing forces. These modes were related to phases of the task, including the onset, offset, holding periods, as well as phases of increasing and decreasing force. These results will inform the design of intracortical brain-computer interface (iBCI) systems that can leverage these naturally occurring patterns of grasp and force control to restore dexterous hand function.

**Significance Statement:** Restoring dexterous hand function after injury remains a major challenge, partly due to an incomplete understanding of the cortical dynamics underlying grasping and force control. In this study, we investigated neural activity within the motor and somatosensory cortices of individuals with tetraplegia attempting to perform grasps to different target forces with varying temporal profiles. We identified distinct neural modes modulated during specific phases of grasp that encode force information throughout the task. These findings suggest that brain-computer interfaces could leverage these native neural modes to restore grasping and force modulation.

## Introduction

Restoration of upper limb motor function is a clinical priority for individuals with cervical spinal cord injury (Anderson, 2004; Simpson et al., 2012; Collinger et al., 2013a; Thorogood et al., 2023). Intracortical brain-computer interfaces (iBCIs) aim to restore arm and hand function by decoding motor intention from the motor cortex, enable intuitive effector control over multiple degrees of freedom (Hochberg et al., 2012; Collinger et al., 2013b; Wodlinger et al., 2015; Bouton et al., 2016; Ajiboye et al., 2017; Guan et al., 2023). However, achieving dexterous object manipulation, which requires precise force control, remains a major challenge for iBCI systems (Pandarinath and Bensmaia, 2022). This limitation partly arises from an incomplete understanding of the cortical dynamics underlying grasp and force control within the sensorimotor cortex.

Force- and grasp-related activity has been observed in the motor cortex (MC) in both humans (Downey et al., 2018; Rastogi et al., 2020, 2021; Okorokova et al., 2024) and non-human primates (Evarts, 1968; Smith et al., 1975; Hepp-Reymond et al., 1978; Cheney and Fetz, 1980; Maier et al., 1993; Oby et al., 2013; Mollazadeh et al., 2014; Menz et al., 2015; Rouse and Schieber, 2016; Okorokova et al., 2024). Early intracortical recordings revealed diverse force-tuning properties at the single-neuron level, where unit activity closely correlates with muscle activity and force output (Evarts, 1968, 1969; Smith et al., 1975; Wannier et al., 1991). At the population level, ensembles of MC neurons are flexibly recruited across diverse motor behaviors, with populations encoding force often overlapping those representing grasp type or other kinematic features. The extent of this overlap remains debated: some studies suggest relative independence of force information (Hendrix et al., 2009; Moreno et al., 2025), while others report interaction between force and kinematic parameters (Degenhart et al., 2011; Intveld et al., 2018; Rastogi et al., 2021). Such overlap may limit the utility of force-related signals during complex behaviors, as evidenced by reduced grasp information during simultaneous movement of the arm during object transport (Downey et al., 2018). Advancing dexterous iBCI control thus requires understanding population-level MC dynamics across varied force conditions, including both static and dynamic force applications.

The MC operates within an interconnected sensorimotor network. The primary somatosensory cortex (SC), for example, is also highly active during grasp (Cavina-Pratesi et al., 2010). While much of this activity reflects peripheral sensory input, cortico-cortical communication from MC to SC also contribute, consistent with their dense bidirectional connectivity (Huber et al., 2017; Umeda et al., 2019; Rolls et al., 2022). Under the “forward model” framework (Wolpert et al., 1995), MC sends an efference copy of motor commands to SC, which modulates sensory input to distinguish self-generated from external stimuli (Straka et al., 2018). A landmark study showed that subpopulations of SC neurons in monkeys could differentiate between active and passive reaching, with some firing prior to movement onset, both excitatory and suppressive efference copy signals from MC (London and Miller, 2012). Furthermore, kinematic information can be decoded from SC during imagined movements, even without peripheral input (Jafari et al., 2020). Yet, efference copy signaling related to force application remains poorly understood, complicated by concurrent proprioceptive and tactile feedback during natural motor actions.

Given this complexity, the present study aimed to characterize neural population activity in MC and SC during attempted grasping and to identify the types of information encoded. We employed a population analysis approach that lends explanation to the diversity of results previously reported in single-neuron studies. Two individuals with tetraplegia resulting from cervical spinal cord injury attempted isometric grasps at varying force targets and rates of force application while neural activity was recorded from MC and SC using intracortical microelectrode arrays. We identified distinct neural modes in both regions that encoded specific phase- and force-related information. These findings advance understanding of sensorimotor population dynamics and have implications for developing iBCIs capable of restoring dexterous functional grasp in individuals with paralysis.

## Materials and Methods

### Experimental Design

Data were collected as part of an ongoing clinical trial (NCT01894802) of an intracortical sensorimotor brain-computer interface (iBCI) conducted under an Investigational Device Exemption (IDE) granted by the US Food and Drug Administration and approved by the University of Pittsburgh Institutional Review Board. Two individuals (P2 and P3) with cervical spinal cord injury (SCI) participated in the study and provided informed consent prior to any experimental procedures. For each participant, two intracortical microelectrode arrays (Blackrock Microsystems, Salt Lake City, UT) were implanted in the hand and arm areas of motor cortex on the surface of the precentral gyrus, which we refer to here as MC (Glasser et al., 2016; Willett et al., 2020) (P2: two 88-channel arrays; P3: two 96 channel arrays). Two additional, 32 channel microelectrode arrays were implanted in the hand area of somatosensory cortex (SC) (Fig. 1A). P2 was a 28-year-old male (at the time of implant) with tetraplegia caused by a C5 motor / C6 sensory incomplete ASIA B spinal cord injury that was sustained about 10 years prior to microelectrode implantation surgery. Data were collected 8-9 years after implant. P3 was a 28-year-old male (at the time of implant) with tetraplegia caused by a C6 ASIA B spinal cord injury that was sustained 12 years before implantation surgery. Data were collected approximately 4 years after implant. Both participants have some residual upper arm and wrist motor function, but no hand-related motor function, and therefore cannot execute grasps. Instead, all grasps were attempted by participants. Both participants also have some residual sensation in their hands.

**Figure 1.**
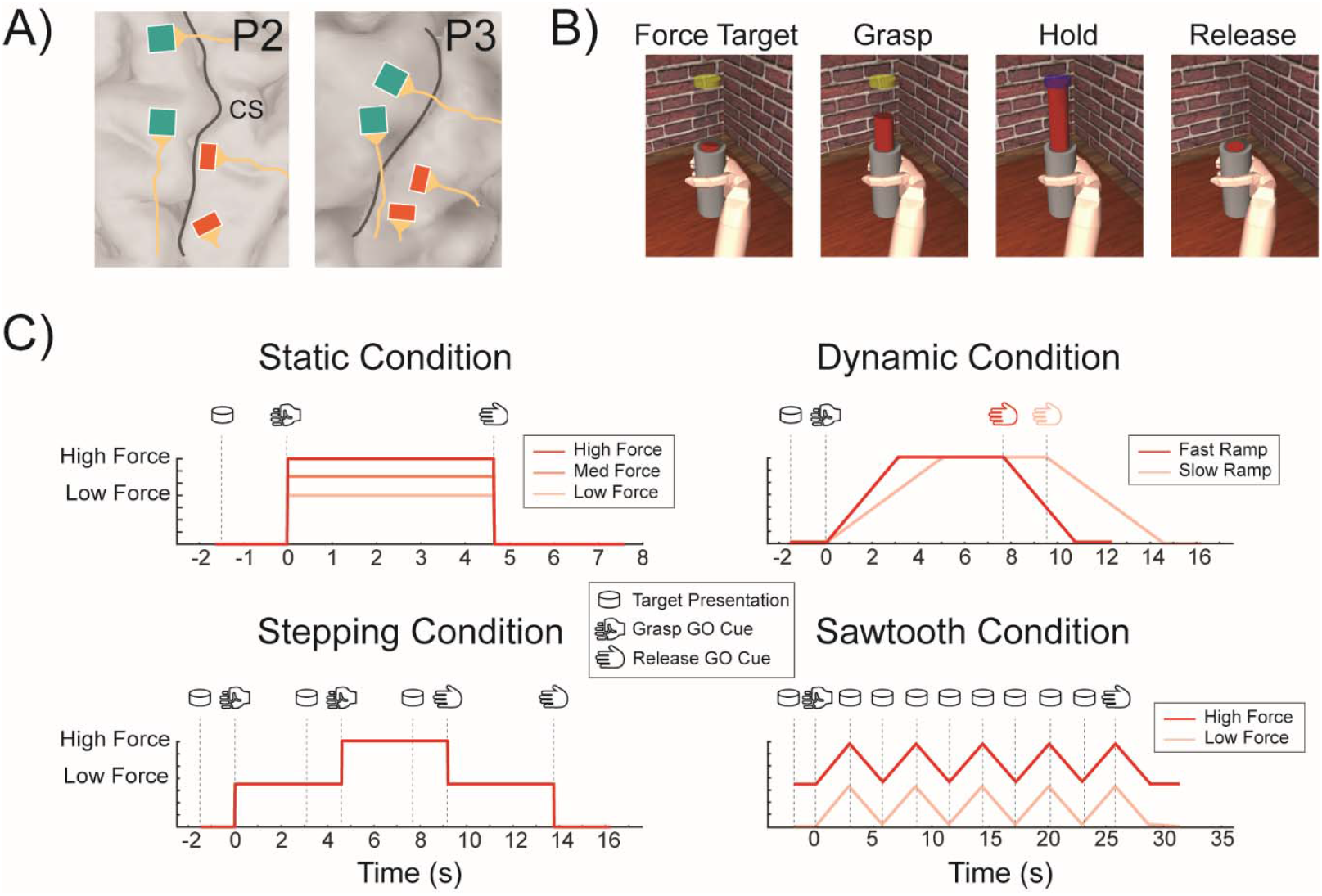
Experimental Design: A) Location and orientation of the MC (green) and SC (red) arrays in the left motor and somatosensory cortex. CS is the location of the central sulcus. B) Depiction of a virtual grasping task at each phase, including a force target presentation phase, a grasp phase, force holding phase, and a release phase. C) Examples of force profiles and task timing for four different grasping conditions. Static conditions consisted of an instantaneous grasp for a target force, a holding period, and an instantaneous release. Dynamic conditions included a ramping phase over a variable time period to a target force, a holding period, and a corresponding ramping down phase to release. More complex conditions included a static force stepping condition, and a dynamic sawtooth condition across two different force ranges.

During an experimental session, participants remained in a position such that the weight of their right arm was supported against gravity. A virtual environment built within the MuJoCo physics simulator (https://mujoco.org/) was displayed on a screen in front of them and, along with audio cues, used to prompt participants to the timing of the task (Fig. 1B). All kinematic and force applications in the simulator were predefined and controlled automatically without participant input or control. In other words, participants were instructed to “follow along” with the actions depicted in the simulation.

Whenever force was applied to the grey cylinder, a smaller red cylinder would rise out of it to a height that represented the currently applied force. All tasks consisted of 1) A force target presentation phase: The force target was represented by a yellow disc placed above the grey cylinder at a specific height for 1.5 s. 2) A grasp phase initiated by an audio GO cue: Virtual force was applied to the grey cylinder and the red cylinder would rise to a height representing the force applied at any given time. 3) A hold phase: When the target force was reached, the force target disk would turn blue. After the specified length of time, the target-force disk would either move to a new height (indicating a new target force) or disappear (indicating that the object should be released) 1.5 s before another audio cue, which cued the 4) release phase, when the participants would release their grasp.

Four different grasping conditions were sampled – static, dynamic, stepping, and sawtooth (Fig. 1C). Static grasping conditions consisted of an instantaneous grasp to a specific force level (high, medium, or low), a period of maintaining, or holding, the target force, and an instantaneous release. To help participants conceptualize force levels, we verbally described the low-, medium-, and high-force targets as equivalent to using the force required to grasp and hold a tomato, orange, and a can of soup respectively, without the object slipping (Downey et al., 2018). Dynamic grasping conditions consisted of force ramping over a set period (1.5 to 6 s) until a high force target was reached, a holding period, and a corresponding force ramping down until full release. Constraining the ramp to variable time windows effectively produced different conditions where the rate of ramp was varied. The stepping condition combined sequential static force grasps within the same trial, specifically, a low-force static grasp, followed by a high-force static grasp, a final low-force static grasp, and release. Finally, the sawtooth condition combined several sequential ramping periods without holding periods. During each session, at least 20 trials of each condition were collected, and at least 6 sessions were carried out per participant.

### Electrophysiology and Data Recording

Neural data was collected from MC and SC via two percutaneous pedestal connectors using NeuroPlex-E headstages connected to two synced Neural Signal Processors (Blackrock Microsystems, Salt Lake City, UT). Signals were hardware filtered between 0.3 – 7500 Hz, sampled at 30,000 Hz, and digitally filtered with a 1^st^ order 750 Hz Butterworth high-pass filter. Spikes exceeding a threshold of −4.5 root mean square (RMS) of the signal on each channel were counted and binned into 20-ms bins.

### Analyses and Statistics

Neural data analysis was performed separately for recordings from MC and SC. Spike counts were square root transformed and convolved with a non-causal Gaussian kernel (σ = 200 ms) to obtain a smoothed estimate of firing rates for each channel. First, we examined responses on individual channels and then we characterized the neural dynamics observed at a population level.

To analyze responses on single channels, all data were grouped by force and rate of force conditions. Every trial was time-aligned within condition to the onset of grasp. For each channel, spike rates were then averaged across trials within condition. Low-firing channels (< 1Hz) were labeled ‘Inactive’ and removed from any further single-channel analyses. Upon visual inspection, we observed different channel specific temporal profiles in the trial-averaged firing rates, including transient activations during onset and offset of grasp as well as tonic activity during static force periods. To classify channels into different response types, we first performed factor analysis (FA) on the averaged spike rates, with time as features and channels as observations. To reduce the number of time bins (∼500–1000 per trial) while minimizing temporal dependence, we divided the trial into non-overlapping 1-s windows and averaged the spike rates within each window. This approach effectively downsampled the data to ∼10–20 features per trial, depending on trial duration.

K-means clustering was then performed on the three factors which explained the most variance of the data. A reasonable number of clusters to use was determined by finding the inflection point of the curve generated when calculating the combined sum of the within-cluster sum of point-to-centroid distances across 1-10 clusters. We used 5 clusters for both P2 and P3 for MC data, 2 for P2 SC data, and 3 for P3 SC data. The channels associated with each cluster were labeled as a ‘class’ according to their shared qualitative firing patterns. Additionally, all channels were visually examined. Some channels that were on the border of 2 or more clusters in factor space were manually assigned to a different class if it was determined that a channel was more representative of that class across all conditions. This process was performed using data from 3 different sessions per participant, treating each channel within a session as an independent observation. Thus, our analysis captures the distribution of observed channel response types across sessions, rather than the activity of longitudinally tracked channels. Finally, the proportion of channels within each class was calculated and compared between participants using a Chi-square contingency table.

For each session, we first estimated the dimensionality of the neural population activity using four-fold cross-validated factor analysis (FA) (Sadtler et al., 2014). For each tested dimensionality (1–30), an FA model was trained on 75% of the data and its log-likelihood evaluated on the remaining 25%. This process was repeated across folds, and the dimensionality that maximized cross-validated log-likelihood was taken as the session’s estimated dimensionality. These results informed our choice to set the latent dimensionality at 20 for subsequent analyses, which captured the majority of explainable variance across sessions while allowing consistency across participants. We then applied FA with 20 factors to the trial-averaged data from each session, followed by a varimax rotation to improve interpretability of temporal features. The rotation was performed on the baseline-centered factors, where the baseline was defined as the averaged response in a 200 ms window beginning 1.5s before the onset of grasp, and the resulting components were then ranked by decreasing explained variance (Dekleva et al., 2024). Finally, to align latent spaces across multiple sessions for each participant, we applied generalized Procrustes analysis (GPA), which finds a linear transformation that minimizes differences between session-specific latent trajectories and a consensus configuration (Gower, 1975). The resulting aligned factors (neural modes) represent dominant, interpretable patterns of population covariation (Gallego et al., 2017). Modes were then labeled by identifying which ones were most strongly modulated during specific phases of the grasp. For single-trial analyses, we projected each trial’s factor scores into the same rotated latent space used for condition averages. Specifically, the orthogonal varimax rotation matrix Q, estimated from the condition-averaged factor trajectories, was applied to the factor time series of each individual trial, resulting in single-trial trajectories in a common factor basis. For each trial and mode we computed the moving sum of squared factor scores within a fixed 200ms window and divided by that mode’s total per-trial variance. We then averaged these time courses across trials (by condition) and across sessions after GPA alignment. A mode’s label (onset, hold, offset, increase, decrease) was assigned by the phase window in which its condition-averaged variance fraction attained its maximum. Because windows overlap, values are not additive across time; we report peaks and phase-integrated fractions for interpretation.

To understand timing relationships between modes, we quantified the latencies between transient peaks and the time points of when relevant modes reached 25, 50, 75, and 90% of their peak modulation during each trial. Mode activations were normalized per trial to their own maxima so that percent thresholds were comparable across trials. Mode-specific search windows were centered around onset and release GO cues +/− 1.5s to identify transient peaks, force holding, and force ramping windows. All landmarks were restricted to these windows. We calculated the timing between transient and holding and ramping related mode landmarks such that the sign convention yields positive values when the mode modulation landmark occurs after the transient. To analyze differences in the timing of neural mode modulation across force conditions (i.e., low vs. high) and modulation levels (i.e., 25, 50, 75, 90%), we performed a 2-way ANOVA on the data to look for main effects, and a post-hoc Šidák correction to control for multiple comparisons family-wise error.

To gain an understanding of force representation in MC and SC, we fit a naïve Bayes classifier (Downey et al., 2018) to each time point across static force conditions using 2 and 3 force levels, and dynamic conditions (without a static component) using 2 force levels. A new model was fit to every classification window, defined using a sliding window of 200ms centered on each time point. GPA-aligned data was used as input, averaged across time within each window. Each classification window was leave-one-out cross validated, and the overall percent correct prediction was calculated. This resulted in a condition-classification curve across the trials. Classification ability is a metric that we defined as the area under this curve bounded by chance levels and relevant phase time points.

To investigate encoding of force rate in the cortex, we performed classification of 4 ramping conditions (1.5, 3.0, 4.5, 6.0 s ramps to a high-force target) using linear discriminate analysis (LDA). We slid a fixed-length window (250 ms, step 50 ms) over each trial. For every window we computed: the mean latent factor vector (across time within the window), and the mean force within the window. The mean force was quantized into bins of width 0.1. To ensure strict force matching and to avoid multiple correlated samples per trial, we enforced one window per trial per force bin. Each retained window was labeled by the ramp duration (4-way classification; one class per instructed ramp duration). LDA was performed with uniform priors, separately for ramp-up and ramp-down windows. 5-fold cross-validation (CV) was performed and was group-aware: folds partitioned trials, and all windows from a trial/group appeared in exactly one fold. Within each fold, features were z-scored using training statistics and any zero-variance columns were dropped. For each force bin we pooled predictions from all CV test folds to obtain a single proportion correct (chance = 0.25). A bin-wise accuracy curve was calculated with a Wilson 95% binomial confidence interval. Bin-wise permutation p-values were computed by repeating the grouped CV after shuffling duration labels within CV groups (thus preserving force matching, window counts, and train/test splits). The p-value is the exceedance probability of the observed accuracy under the null (5,000 permutations). Because multiple bins are tested, we applied Benjamini–Hochberg FDR correction (q=0.05) separately for ramp-up and ramp-down panels.

## Results

### Single channels are tuned to specific phases of the grasp

In studying single channel responses during grasping behavior, we observed channel tuning to specific phases of the grasp in both MC and SC. Across three sessions from each participant (Total Channels: P2: MC = 528, SC = 192; P3: MC = 576, SC = 192), spike rates for a given channel were averaged across all trials within each condition, and K-means clustering was used to group channels with similar firing patterns (Fig. 2A&B). As channel response patterns cannot be stably tracked across sessions, we analyzed each channel/session independently. Reported proportions therefore reflect the distribution of observed response classes across three sessions per participant, not unique neurons. In MC, upon visual inspection, these distinct firing patterns represented transient responses during increases in force, transient responses during decreases in force, transient responses during any change in force (‘multi-transient’), tonic responses during holding periods of constant force, or untuned responses (Fig. 2C). Similar, but fewer, classes were identified in SC, including transient responses during increases in force and transient responses during decreases in force (Fig. 2D). We did not identify any SC channels with the multi-transient or tonic hold responses that were identified in MC, however, the channels with transient responses during increases in force tended to have a longer tail that extended into the holding periods (Fig. 2D).

**Figure 2.**
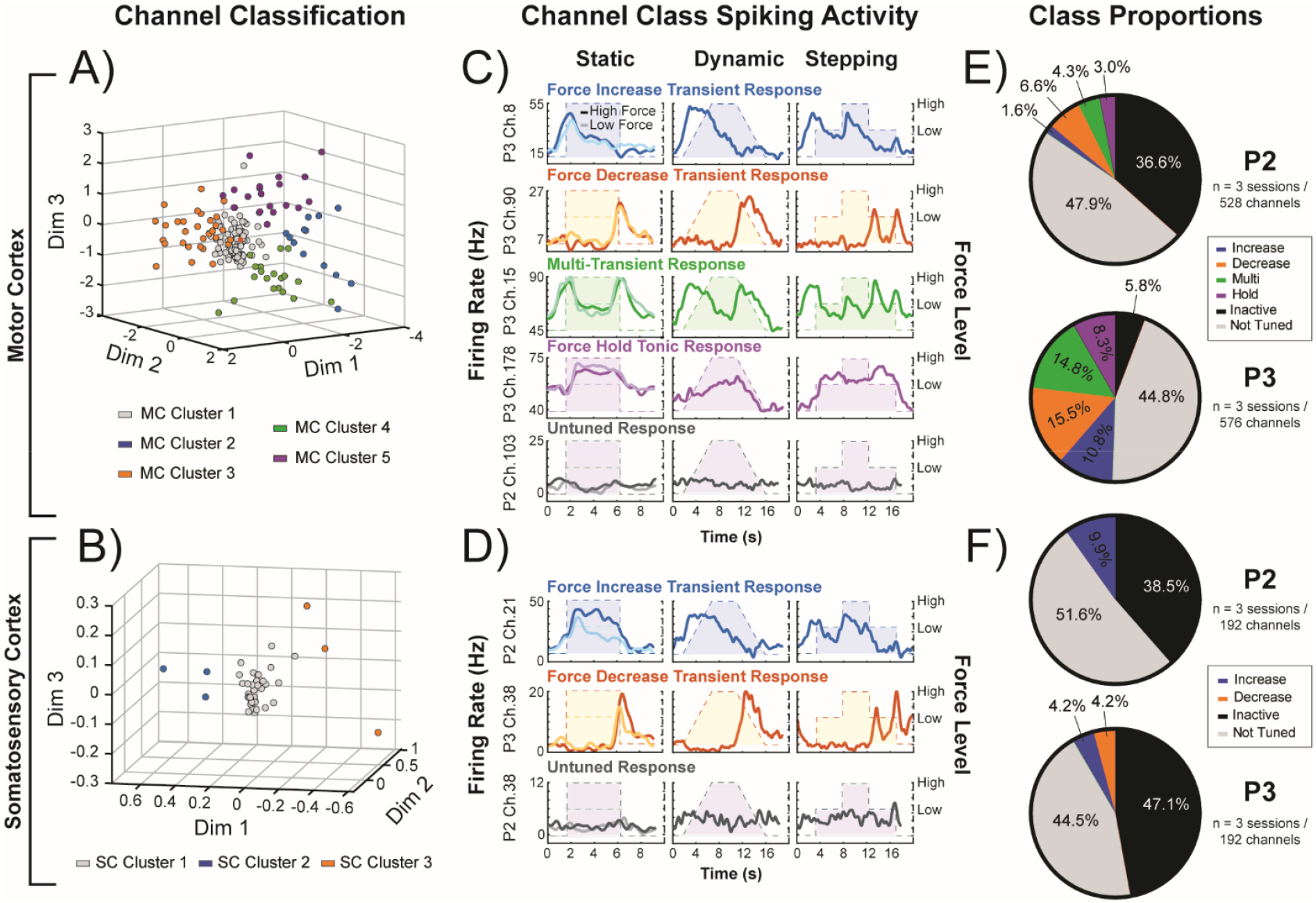
Spiking activity patterns on single channels: A-B) K-means clustering of channel spiking activity during all grasp conditions in A) motor cortex (MC) and B) somatosensory cortex (SC). Each dot represents a channel. An example session from P3 is shown here. C-D) Example channels of trial-averaged responses from each cluster across each grasping condition in C) motor cortex and D) somatosensory cortex. Dotted lines represent the force profile in each condition. Classes were defined for each cluster describing their response profile. Example classes from both participants (P2 & P3) are shown here. E-F) Proportion of channels in each class for E) motor cortex and F) somatosensory cortex.

In P2, the total proportion of MC channels with modulated firing patterns was 15.5%, with each class accounting for between 1.6-6.6% of total channels. In P3, this proportion was 55.2%, with each class accounting for between 8.3-15.5% of total channels (Fig. 2E). This difference is likely due to the durations of the implants with fewer channels in P2 recording neural signals at more than 8 years post-implant. Excluding channels considered inactive and not tuned, we found no significant difference in proportions of classes between participants (*X*^2^ (3, n=361) = 7.43, p = 0.059), suggesting a similar distribution of tuned channel classes across participants. Overall, fewer SC channels were modulated to the task (Fig. 2F). Because we did not identify any decrease-specific channels in SC for P2, no additional statistics were performed on SC class proportions. In summary, we observed that both MC and SC contained channels with transient responses tuned to increases and decreases in force. However, only MC contained multi-transient responses modulated by both increases and decreases in force, as well as tonic responses during the hold phase. These results indicate that, while both MC and SC demonstrate multi-unit task related tuning, MC exhibits a broader repertoire of force-related response patterns than SC, consistent with its primary role in initiating and sustaining grasp forces.

### Distinct MC neural modes capture phase-specific activity during grasping

Having demonstrated the variety of individual channel responses, we went on to analyze the population response of the sensorimotor cortex during grasping. We did this by examining the dominant, interpretable patterns of population covariance, or neural modes, from both the MC and SC during the attempted behavior (see Methods). Unlike per-channel labeling, population activity can capture coherent modes from groups of weakly modulated neurons and yields orthogonal dimensions that partition shared variance into distinct components (Cunningham et al., 2014). In both participants, individual neural modes were consistently modulated during specific phases of the grasp (e.g., onset, hold, release), and this phase-locked modulation was preserved across all grasping conditions.

In MC, the top 10 neural modes captured approximately 90.4% of the variance explained in P3, and approximately 89.7% of the variance explained in P2, whereas the top 3 neural modes captured approximately 43.4% and 61.3% of the variance in P3 (Supplementary Fig. 1A) and P2 (Supplementary Fig. 1D) respectively. Labels were assigned to modes based on where each mode’s fractional variance peaked relative to task phases (Fig. 3A&E). In MC for P3, we identified 6 task-relevant neural modes, which captured the following responses: a grasp onset-related transient component, a grasp offset-related transient component, a grasp hold-related static component modulated during high forces, a grasp hold-related static component modulated during low forces, a component modulated during increasing forces, and a component modulated during decreasing forces (Fig. 3B).

**Figure 3.**
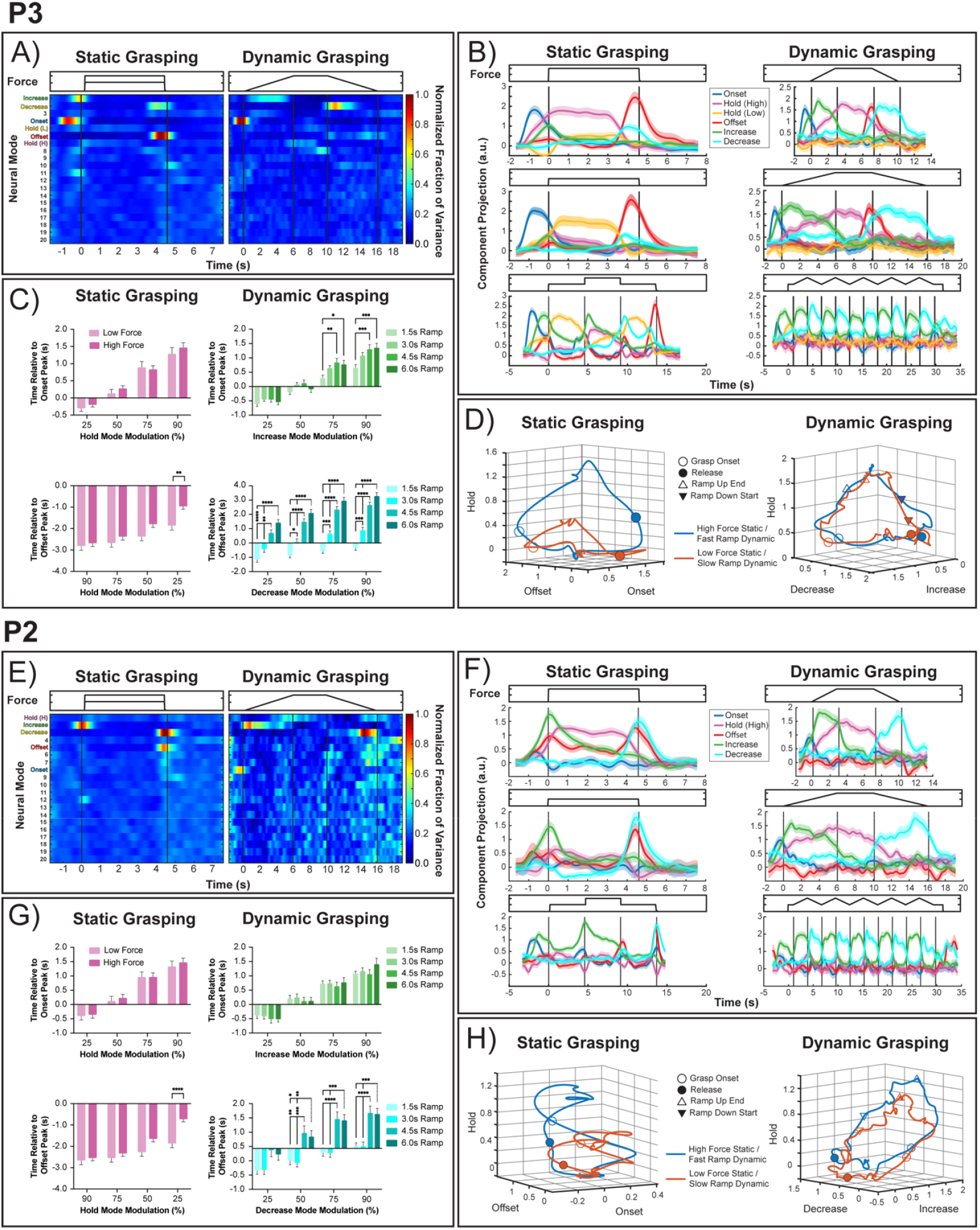
MC Neural Modes for participant P3 (A-D) and P2 (E-H): A&E) Heatmaps show temporal concentration of variance for each mode. Color values are normalized to the global maximum across all modes, such that 1 represents the strongest hotspot observed. Because windows overlap, values across time are not additive. B&F) Trial-averaged neural modes during static and dynamic grasping conditions. Modes represent GPA-aligned data from three sessions per participant. Shaded regions represent the 95% CI across trials (N=60), pooled across 3 sessions after GPA alignment. C&G) Mean latency (+/− SEM) of when the holding-related mode (left) and the increase / decrease modes (right) reached 25-90% of their total modulation relative to the peak of the onset transient (top) and the offset transient (bottom) (**** p < 0.0001, *** p < 0.001, ** p < 0.01, * p < 0.05). Dashed significance bars indicate the group that all other comparisons are made against during post-hoc analysis. D&H) Neural data projected along the onset, offset, and hold (high) dimensions during static grasping (left) and along the increase, decrease, and hold (high) dimensions during dynamic grasping (right).

Largely similar task-relevant modes were observed in P2 (Fig. 3F), but their relative engagement differed from P3. The hold-related static component that showed clear modulation at low forces in P3 was not evident in P2. During static grasping, P2 relied more heavily on the force increase- and decrease-related modes relative to his own onset-related mode, whereas in P3 the onset-related mode was the dominant component during this phase. During dynamic grasping, the offset-related mode in P2 showed only weak deviations from baseline, in contrast to the pronounced offset-related modulation observed in P3.

By comparing the time when the hold and ramp-related modes reached specific modulation levels (25, 50, 75, or 90% of its total modulation) relative to the peaks of the onset and offset transients, we found distinct timing relationships for static versus dynamic grasping (Fig. 3C&G). In both participants, the hold mode reliably followed onset transients (top left panel) and preceded offset transients (bottom left panel) during static grasping. The onset → hold latencies were not different across force levels for either P3 [Force Cond F (1, 472) = 0.9920, p = 0.3198; Mod Level F (3, 472) = 58.88, p < 0.0001; Interaction F (3, 472) = 0.3306, p = 0.8033] or P2 [Force Cond F (1, 672) = 0.4457, p = 0.5046; Mod Level F (3, 672) = 48.41, p < 0.0001; Interaction F (3, 672) = 0.07569, p = 0.9731], indicating that the timing of grasp onset into sustained holding activity in MC is force-invariant. We also looked at the end of the holding period when the grasp transitions from hold to release. An overall timing difference of the hold → offset latencies across force conditions was seen in both P3 [Force Cond F (1, 472) = 12.72, p = 0.0004; Mod Level F (3, 472) = 19.51, p < 0.0001; Interaction F (3, 472) = 1.644, p = 0.1783] and P2 [Force Cond F (1, 664) = 17.42, p < 0.0001; Mod Level F (3, 664) = 21.96, p < 0.0001; Interaction F (3, 664) = 3.510, p = 0.0151] with post hoc analyses revealing a difference only at the 25% level for both participants (bottom left panel). In both cases, the hold component decreased to 25% of its peak value earlier for the low force condition. However, this difference at lower modulation levels may just be due to the low overall modulation of the holding-related mode at low forces.

During dynamic trials, the ramping window, or rate of ramp, was varied across conditions but maintained within each trial until the same force target was achieved across conditions. The timing between transients and ramp-related modes was dependent on ramp duration but showed evidence of a plateau for longer ramps. For P3, both the onset → increase [Ramp Cond F (3, 1088) = 9.45, p < 0.0001; Mod Level F (3, 1088) = 144.2, p < 0.0001; Interaction F (9, 1088) = 1.319, p = 0.2224] (Figure 3C, top right panel) and offset → decrease [Ramp Cond F (3, 1029) = 162.9, p < 0.0001; Mod Level F (3, 1029) = 34.16, p < 0.0001; Interaction F (9, 1029) = 1.165, p = 0.3142] (Figure 3C, bottom right panel) comparisons showed an overall effect of ramp duration, where in general, the increase and decrease modes peak later in time relative to their respective transients as ramp durations increase. For P2 the effect of ramp duration was not significant for onset → increase [Ramp Cond F (3, 944) = 0.6585 p = 0.5777; Mod Level F (3, 944) = 103.4, p < 0.0001; Interaction F (9, 944) = 0.4634, p = 0.8994] (Figure 3G, top right panel), but it did reach significance for offset → decrease [Ramp Cond F (3, 869) = 32.79, p < 0.0001; Mod Level F (3, 869) = 23.33, p < 0.0001; Interaction F (9, 869) = 0.5254, p = 0.8568] (Figure 3G, bottom right panel), where shorter decrease ramps reached modulation thresholds earlier than longer ramps, similar to P3. Post hoc comparisons revealed that these differences were driven primarily by the shorter ramps (1.5 and 3.0s), whereas the relative latencies for the longer ramp conditions (4.5 and 6.0s) were statistically similar. Thus, the increase and decrease modes appear to be rate-dependent during shorter ramps but saturate during longer ramp durations, consistent with a bounded sensitivity of MC dynamics to ramp speed. Together, these analyses demonstrate that MC transient and sustained/ramp modes are coordinated in a task-specific manner: the timing of the “handoff” between transient (onset and offset) and holding-related modes are independent of force level, while the timing of the “handoff” between transient and ramping-related modes scale with rate of force, especially with shorter ramps. However, with longer ramps, ramping-related modes reach maximal modulation at similar time points relative to transient modes, independent of rate.

Projections of the high- and low-force static grasping conditions onto the most prominent neural modes revealed a clear separation along the high force-specific hold-related dimension (Fig. 3D&H). This separation was consistent across both participants and across trials, supporting the idea that absolute force level is robustly encoded in dedicated hold-related modes. Similar force condition separation was observed when projecting the same data along the low force specific hold-related dimension. In contrast, fast and slow (3s and 6s ramp duration) dynamic grasping conditions traced largely overlapping trajectories along the force-increase and force-decrease modes. This overlap suggests that rate of force change was not strongly represented in the low-dimensional MC population dynamics. Taken together, the latency analysis (Fig. 3C&G) indicates that the temporal coordination between onset/offset transients and the increase/decrease modes varies with ramp kinematics, whereas the trajectory plots (Fig. 3D&H)—restricted to the increase–decrease–hold subspace—show broadly overlapping ramp trajectories during the ramp itself, suggesting limited separability of ramp conditions in these dimensions once the ramp is underway.

### SC neural modes are modulated during attempted grasp without afferent input

Despite a lack of somatosensory input during attempted motor behavior, we observed task-related modulation of neural modes during grasping in SC, similar to what we observed in MC. Overall, dimensionality was lower than in M1: the top 3 modes explained ∼66.3% of the variance in P3 and ∼62.0% of the variance in P2 (Supplementary Fig. 2A,D), whereas the top 10 neural modes explained ∼88.0% and ∼86.4% respectively (however, this may be due to fewer recording electrodes in SC – see Methods). Labels were assigned based on fractional variance peaks relative to task phases (Fig. 4A&E). In both participants, three task-relevant neural modes were identified: a grasp onset-related mode, a force-increase mode, and a force-decrease mode (Fig. 4B&F). Compared to MC, the repertoire of SC modes was narrower. There was no clear hold-related mode or offset-related transient. Instead, the increase mode displayed a slow decay that extended into the hold period, suggesting that SC may indirectly represent sustained force levels through prolonged increase-related activity rather than via a dedicated tonic ‘hold’ mode. P3 showed robust modulation of both increase and decrease-related modes, while P2 showed much weaker modulation of the decrease mode during dynamic grasping.

**Figure 4.**
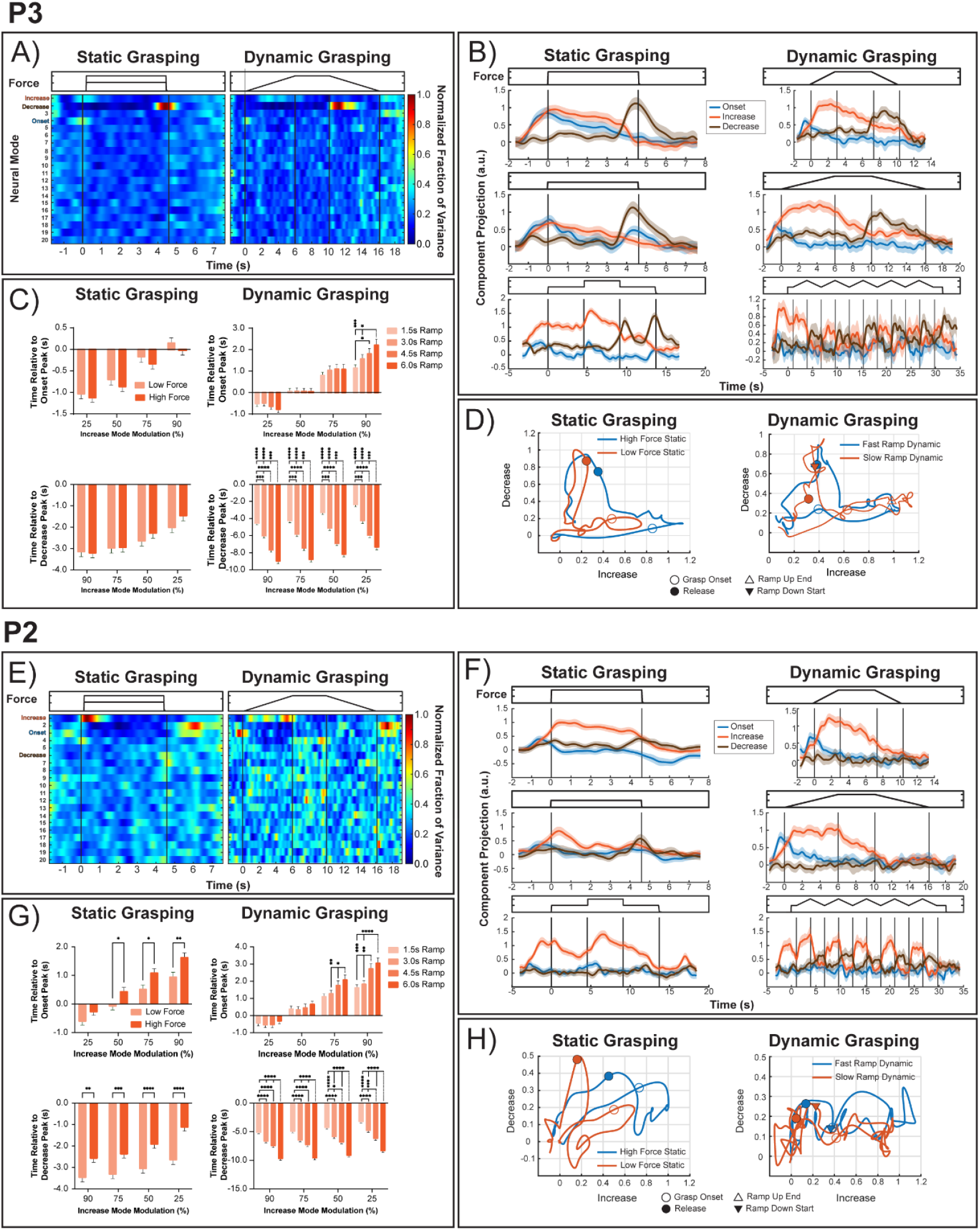
SC Neural Modes: A&E) Heatmaps show temporal concentration of variance for each mode. Color values are normalized to the global maximum across all modes, such that 1 represents the strongest hotspot observed. Because windows overlap, values across time are not additive. B&F) Trial-averaged neural modes during static and dynamic grasping conditions. Modes represent GPA aligned data from three sessions per participant. Shaded regions represent the 95% CI across trials (N=60), pooled across 3 sessions after GPA alignment. C&G) Mean latency (+/− SEM) of when the increase mode reached 25-90% of its total modulation relative to the peak of the onset transient (top) and the decrease mode (bottom) (**** p < 0.0001, *** p < 0.001, ** p < 0.01, * p < 0.05). Dashed significance bars indicate the group that all other comparisons are made against during post-hoc analysis. D&H) Factor projections along the Increase and Decrease dimensions during static (left) and dynamic (right) grasping.

Latency analyses during static grasping compared the onset transient peak to the percent modulation of the increase mode (Fig. 4C&G, top left). In P3, the onset peak and the 90% level of the increase mode occurred close in time, indicating that the increase mode was recruited almost immediately following grasp onset during static grasping. While an overall difference in force condition was observed [Force Cond F (1, 472) = 3.872, p = 0.0497; Mod Level F (3, 472) = 46.12, p < 0.0001; Interaction F (3, 472) = 0.07703, p = 0.9724], post hoc analysis revealed no significant differences between force levels at any modulation level. In P2, by contrast, the increase mode rose more slowly, with the 90% level reached noticeably later (up to 1.5 sec) than the onset peak. Moreover, onset → increase latencies were significantly later for the high force condition compared to low force [Force Cond F (1, 672) = 31.30, p < 0.0001; Mod Level F (3, 672) = 64.32, p < 0.0001; Interaction F (3, 672) = 0.6185, p = 0.6032], suggesting that in P2, phase transitions during static force conditions in SC was partially modulated by force.

We next measured when the increase mode reached 25–90% modulation relative to the peak of the decrease mode (Fig. 4C&G, bottom left). Across participants, the increase mode’s 25–90% landmarks occurred before the decrease peak (i.e., negative latencies), consistent with a slow decay of force increase-related activity that extends into the hold period but subsides ahead of release. In P3, no effect of force was found [Force Cond F (1, 466) = 2.199, p=0.1388; Mod Level F (3, 466) = 18.18, p < 0.0001; Interaction F (3, 466) = 0.9091, p = 0.4365] suggesting that the relative timing of the increase and decrease components was consistent in both force conditions. In P2, however, these relative latencies differed significantly across force levels [Force Cond F (1, 659) = 78.54, p < 0.0001; Mod Level F (3, 659) = 15.61, p < 0.0001; Interaction F (3, 659) = 1.270, p = 0.2838], such that the increase mode in the high force condition was sustained longer throughout the trial and decreased closer to the decrease mode peak than the increase mode in the low force conditions, further supporting force dependence during static grasping in SC for this participant.

For dynamic conditions, we quantified when the increase mode reached 25–90% of its total modulation relative to the onset peak (Fig. 4C&G, top right). In P3, there was no significant effect of ramp duration, but there was a significant interaction between ramp duration and modulation level [Force Cond F (3, 1088) = 1.887, p=0.1301; Mod Level F (3, 1088) = 163.2, p < 0.0001; Interaction F (9, 1088) = 2.400, p = 0.0108]. Post hoc analyses indicated that ramp-related differences emerged only at the 90% level. Shorter ramps (1.5 and 3.0 s) which required faster force application reached late-phase recruitment earlier than longer ramps, whereas 4.5 vs. 6.0 s ramps did not differ, indicating a plateau at longer durations. Thus, in P3, ramp duration influenced the onset → increase relationship only in the late phase of increase-mode recruitment, consistent with a rapid early saturation followed by limited sensitivity to ramp duration. In P2, ramp effects emerged at higher modulation levels (75 and 90%)—late portions of the increase trajectory were delayed for longer ramps—while the 25 and 50% levels showed no reliable separation [Force Cond F (3, 944) = 12.79, p<0.0001; Mod Level F (3, 944) = 177.4, p < 0.0001; Interaction F (9, 944) = 2.645, p = 0.005]. Post-hoc tests again showed that differences were driven by 1.5–3.0 s ramp conditions, with no significant difference between 4.5 and 6.0s. Together, these results suggest that late-phase recruitment of the SC increase mode is rate-dependent at shorter ramps but saturates at longer durations, with P2 showing more graded late-phase sensitivity (75/90%) and P3 showing earlier saturation, with effects only at the 90% level.

When aligned to the decrease peak, the increase mode consistently preceded the decrease mode (Fig. 4C&G, bottom right). In this comparison, we are specifically tracking the decay of the increase mode during the hold period leading up to release. In P3, this decay scaled robustly with ramp duration [Force Cond F (3, 1024) = 331.3, p<0.0001; Mod Level F (3, 1024) = 50.91, p < 0.0001; Interaction F (9, 1024) = 2.836, p = 0.005], with shorter ramps showing faster decline and a plateau between 4.5 and 6.0 s. In P2, despite weaker overall modulation of the decrease mode, the increase → decrease relationship also showed significant ramp effects [Force Cond F (3, 934) = 280.4, p<0.0001; Mod Level F (3, 934) = 35.59, p < 0.0001; Interaction F (9, 934) = 0.4029, p = 0.005], again driven by shorter ramps. Thus, across both participants, the decay of increase activity into release was stretched for longer ramps but plateaued at longer durations, reinforcing that in SC, the increase mode partially substitutes for a missing hold mode by bridging the ramp-up and ramp-down phases.

Factor projections of the static grasping conditions along the most prominent SC modes showed a modest separation along the force-increase dimension, suggesting some encoding of absolute force (Fig. 4D&H, left). This separation was weaker than the clear hold-related separation seen in MC, consistent with the absence of a dedicated hold mode in SC. Instead, the slow decay of the increase mode during static trials may provide a partial surrogate for force representation. In the dynamic condition (Fig. 4D&H, right), P3 trajectories for fast and slow ramps were largely overlapping along the increase and decrease dimensions, providing little evidence for explicit encoding of ramp speed. For P2, interpretation was limited by the minimal modulation of the decrease mode, which reduced the dimensional separation of trajectories. This again contrasts with MC, where ramp timing effects were clearer, and supports the idea that SC contributes primarily to representing absolute force levels rather than force-rate dynamics.

Together with the latency analyses, these projection results reinforce that SC dynamics differ from MC: absolute force is weakly represented through the increase dimension, while ramp trajectories lack strong separation by rate of force application. The increase mode’s decay into the hold and release periods likely accounts for much of the static separation, underscoring its role as a surrogate for the missing hold-related neural mode in SC.

### Motor cortex encodes absolute force during both static and dynamic grasping

We next asked how neural populations in MC and SC encode force across static and dynamic conditions. Force encoding was first evaluated using a naïve Bayes classifier trained to discriminate force levels within each condition. In MC, classification accuracy for both 2- and 3-force levels was consistently above chance (Fig. 5A&B), peaking early in the hold phase and remaining elevated throughout much of the grasp before declining near offset. In SC, accuracies were lower than MC, peaking close to grasp onset for both participants in the 2-force condition. For the 3-force condition, peak accuracy occurred near grasp onset in P3 but later in the hold period in P2. These results indicate that both MC and SC carry force information during static grasps, with stronger and more sustained encoding in MC.

**Figure 5.**
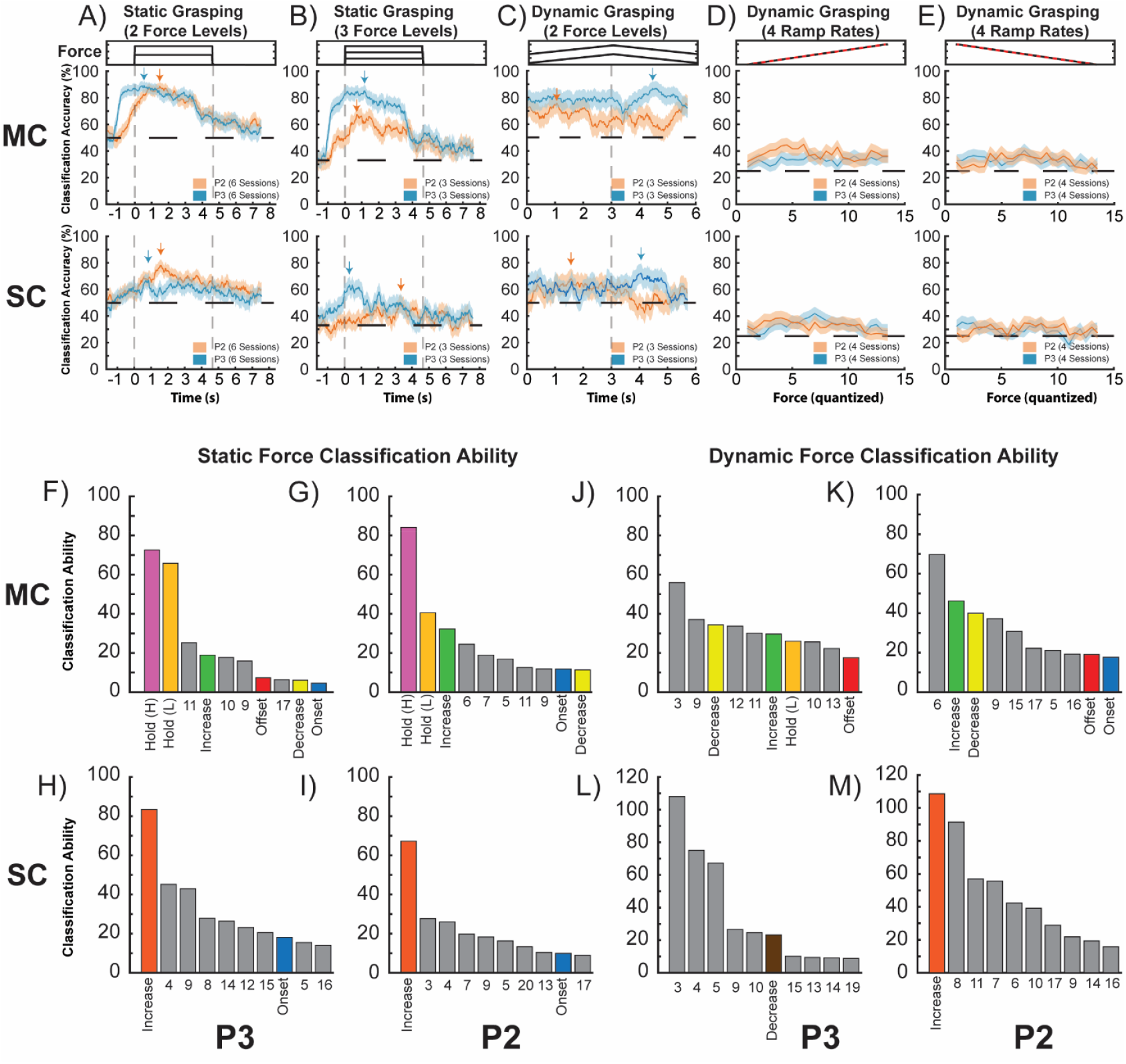
Force classification: A-C) Force classification accuracies at each time point during static grasping with 2 force levels, static grasping with 3 force levels, and dynamic grasping (2 force levels) in both MC and SC Arrows indicate the peak classification accuracy. Solid lines depict the mean classification accuracies and the shaded regions represent the variance over 10,000 bootstraps. D&E) Force ramping rate classification with 4 different ramps compared across force aligned windows. F-M) Classification abilities of individual neural modes in MC and SC for both participants. F-I) results are for a 2-force static grasping task similar to the classification results with all neural modes shown in A. J-M) results are for a dynamic grasping condition with two force levels similar to the classification results with all neural modes shown in C.

For dynamic grasping, we classified force conditions during the averaged dynamic periods of the sawtooth condition (i.e., ramp up and ramp down, Fig. 5C). Classification accuracies remained above chance level in MC for both participants throughout both the ramp up and ramp down phases. This peaked during the ramp up phase for P2, and during the ramp down phase for P3. However, the overall range of accuracies throughout the condition were lower than the static conditions. Like the static conditions, classification accuracies for the dynamic condition in SC were lower than in MC, and they remained relatively stable throughout the grasp for both participants.

We also asked whether neural activity distinguishes ramp rate (1.5, 3.0, 4.5, 6.0 s ramps to the same high force target) after controlling for absolute force. Rate decoding accuracy was slightly above chance for both participants and for both directions (Fig. 5D: ramp-up; Fig. 5E: ramp-down). Accuracy varied slowly with force, indicating that modest rate information is present across the force range rather than being confined to a specific level. Few individual bins reached significance—unsurprising given the modest per-bin effect sizes and multiple comparisons—but the aggregate pattern (median accuracy across bins and area-over-chance) was positive and similar across participants. These results show that while MC and SC contain some rate-related information, absolute force encoding appears to be favored.

To understand how each individual neural mode contributes to force encoding throughout the grasp, we re-trained and tested a Naïve Bayes classifier using each neural mode as a single feature. We then calculated the classification ability of each mode by finding the area under the classification accuracy curve bound by the chance level on the y-axis and the relevant behavioral phase windows on the x-axis. This area was taken as a proportion of the area calculated when classifying using the full feature set. Effectively, classification ability is a measure of how well an individual neural mode can classify grasping force conditions. In MC during static grasping (2-force levels), the hold (high), hold (low), and force increase related neural modes had notable overall classification abilities for both participants: 72.7%, 65.8%, and 18.9% respectively for P3 and 84.1%, 40.5%, and 32.2% for P2 (Fig. 5F&G). In SC, the force increase mode had the highest overall classification ability for both participants: 83.4% for P3 and 67.2% for P2, which was most effective at classifying force during the hold phase of the grasp (Fig. 5H&I).

During dynamic grasping (2-force levels), unlabeled MC modes #3 and #9 (Supplemental Fig. 1C) had the greatest classification ability for P3 (Fig. 5J). The modulation patterns of these modes were similar to that of the force decrease and offset modes during the low-force dynamic condition but were relatively inactive during high-force dynamic as well as static conditions, explaining their high classification abilities. Similarly, for P2, neural mode #6 (Supplemental Fig. 1E) which had a modulation pattern resembling that of the offset mode only during the low-force dynamic condition, had the highest classification ability (Fig. 5K). This was followed by the force-increase and force-decrease modes. In SC during dynamic conditions, overall classification accuracies were low for both participants (Fig 5C). For P3, neural modes #3-5 had the highest classification abilities but had inconsistent activation patterns across grasping conditions (Fig. 5L). For P2, the force increase mode had the largest classification ability (Fig. 5M).

Overall, during static grasp for both participants the holding-related and force increase-related modes were most informative of force in MC, and the force increase-related mode was most informative of force in SC. During dynamic grasp, results were more varied. The force-increase and -decrease related neural modes were informative of force in MC for both participants, as was the force-increase related mode in SC for P2.

## Discussion

### Summary of Main Findings

This study investigated cortical dynamics in motor (MC) and somatosensory cortex (SC) during static and dynamic grasp force tasks in participants with tetraplegia. We found that even without overt movement, attempted grasping robustly engaged MC neurons and modulated SC neurons in a task-relevant manner. At the single-channel level, MC exhibited transient and tonic responses tuned to grasp phases, while SC showed similar but less sharply tuned activity. At the population level, distinct and separable neural modes emerged within each cortical area, reflecting unique functional dynamics. In MC, modes were modulated by grasp phase and informative of force condition (low vs. high force), with holding-related modes contributing most to force encoding. SC modes were also tuned to task phase and, to a lesser degree, to force condition. Latency analyses and classification of force rate indicated that both MC and SC preferentially encoded absolute force rather than rate of force change.

### Response characteristics on the single channel level

Single-channel analyses revealed distinct firing patterns—transient and tonic—related to attempted grasping in MC, consistent with findings from intact primate motor systems performing precision grasps. Similar subpopulations displaying phasic, tonic, ramping, or combined phasic–tonic patterns have been observed across the motor cortex (Smith et al., 1975; Hepp-Reymond et al., 1978; Wannier et al., 1991; Maier et al., 1993; Hendrix et al., 2009), cerebellum (Smith and Bourbonnais, 1981), motor thalamus, globus pallidus (Anner-Baratti et al., 1986), as well as in spinal premotor interneurons (Takei and Seki, 2013), supporting consistency of force-encoding motifs across the neuroaxis.

We extend these results by showing similar firing patterns in the MC of individuals with paralysis during attempted grasp. Unlike earlier work, we did not observe neurons whose firing decreased with force. This may reflect the grasp type used: inhibitory responses appear more prominent during dexterous grasps requiring fine control (Wannier et al., 1991). Participants here performed power grasps, which may engage broader excitatory populations with less cortical inhibition. Indeed, grasp type interacts with force encoding across overlapping populations (Intveld et al., 2018; Rastogi et al., 2021).

Interestingly, phasic–tonic responses were less common in our dataset. A likely explanation is the absence of afferent feedback. MC activity is known to be modulated by tactile input (Tokimura et al., 2000; Lei and Perez, 2017; Bao et al., 2024), and by intracortical microstimulation (ICMS) of SC (Osborn et al., 2021; Shelchkova et al., 2023). Combined phasic–tonic activity in intact systems may therefore reflect ongoing sensorimotor communication between SC and MC—a mechanism attenuated here due to loss of peripheral input.

SC, while classically viewed as sensory-driven, is also active during motor planning (Gale et al., 2021; Ariani et al., 2022), observation (Kuehn et al., 2014), and imagery (Jafari et al., 2020; Wandelt et al., 2022). fMRI studies have similarly shown that attempted finger movements in individuals with tetraplegia activate preserved SC representations (Kikkert et al., 2021; Downey et al., 2024). Our spiking-level data extend this evidence, revealing task-related modulation of SC neurons during attempted grasp. Although firing rates and the proportion of tuned channels were lower in SC than in MC, similar transient activity patterns were evident.

Given the absence of somatosensory input, SC activity may be explained by efference copy signaling - internal motor command signals that help distinguish reafferent from external sensory events (Straka et al., 2018). Notably, we did not observe the tonic SC responses typical of intact animals (Wannier et al., 1991), but rather elongated transient activity that extended into the holding phase. This pattern may reflect sustained efferent signaling from MC during force increases or maintenance. In intact systems, such signals might be masked by strong peripheral feedback, highlighting the advantage of the current paradigm for isolating cortical communication.

### Distinct neural modes in both MC and SC

Population-level analyses revealed that MC activity decomposed into multiple distinct neural modes corresponding to functional task components. Transient modes were engaged during rapid force changes (grasp onset and release), while tonic modes dominated during slower force modulation and sustained holds. Similar decompositions of MC activity into transient and tonic components have been observed during both overt and imagined isometric wrist movements (Dekleva et al., 2024). Here, we expand on those findings by showing that such neural modes also underlie attempted static and dynamic grasping.

These modes suggests that MC flexibly partitions its population activity into distinct functional subspaces, supporting independent control of dynamic versus static force components. While individual channels sometimes resembled these motifs, the population approach provided a richer view. Neurons with mixed selectivity (e.g., transient plus weak tonic responses) were naturally captured through shared loadings across multiple modes, whereas single-channel analyses may misclassify or overlook such structure. Factor analysis (FA) further isolates orthogonal components of variance, yielding interpretable, nonredundant subspaces—an advantage over categorical neuron-level classification. Thus, neural modes provide a complementary perspective: they recapitulate familiar response motifs while quantifying the underlying population-level covariance structure.

The presence of dedicated holding-related modes indicate that MC contributes to sustaining force, not just generating transients. Yet, consistent with prior evidence, the population encoding of dynamic activity was broader and stronger than that for steady output. Noninvasive studies (EEG, ECoG, fMRI) similarly report bursts of MC activity during dynamic force production but limited sustained activation during static hold (Thickbroom et al., 1999; Keisker et al., 2010; Flint et al., 2014; Jiang et al., 2020). Despite this, we identified robust MC activity during holding phases, and these modes carried the most information about absolute force.

Given that MC populations largely encode overlapping motor features (kinetic and kinematic), it remains unclear how “resilient” holding-related neural modes are to added motor complexities, such as during grasp-and-carry behavior. Previous iBCI work from our lab demonstrates that force decoding throughout the holding period of a task such as this is difficult (Downey et al., 2018), while decoding the transient signals during dynamic phases such as grasp onset and offset can be robust (Dekleva and Collinger, 2025). Unlike previous studies (Smith et al., 1975; Hepp-Reymond et al., 1978), we did not find that MC encoded for rate of force change, however, we were still able to classify absolute force throughout dynamic ramping periods. MC activity during these phases was primarily composed of force-increase or force-decrease related neural modes, which also carried force-related information during dynamic phases of grasp.

One limitation of studying attempted movements is the reliance on cues to determine the timing of phases for behavior. It is highly unlikely that the participants initiated grasp onset and release exactly when they were cued to do so. This can explain some of the variable latencies that we observed across conditions in our neural mode analysis and during force classification. For example, it appears that P3 tended to both initiate and release grasps earlier than cued. Further, our task design included a fixed 1.5 s target presentation period before the GO cue, which likely contributed to anticipatory activity in motor cortex during the pre-grasp period, as participants could reliably predict the onset of the movement cue. Such anticipatory modulation should be considered when interpreting early neural responses.

### Neural Modes in Somatosensory Cortex

Notably, we found distinct neural modes in SC during grasp and force modulation. Overall dimensionality in SC was lower than in MC, which was unsurprising given the lack of afferent input during the task. Three neural modes were consistently modulated across sessions and conditions, independently associated with the onset of grasp, increases in force, and decreases in force. However, P2 displayed relatively weak modulation of the decrease related mode, which was not a surprising result given the lack of classified force decrease transient channels in SC for that participant. Even in SC, we were able to classify force conditions above chance levels during both static and dynamic conditions in both participants. Overall classification accuracies were lower in SC than in MC, however, this was not unexpected given that we had fewer overall recording channels and fewer tuned channels in SC.

In conclusion, our results demonstrate that even in the absence of movement and somatosensory feedback, the human sensorimotor cortex retains rich representations of both transient (dynamic) and tonic (static) phases of grasp force. MC and SC each engage distinct neural modes to handle changes in force versus holding steady forces, indicating partially independent control schemes for dynamic versus static force regulation. These insights advance our understanding of how the brain orchestrates complex hand actions and may inform the development of assistive technologies. In particular, iBCI systems for grasp restoration could leverage both the transient and tonic signals related to force change and maintenance to achieve more natural and effective control of neuroprosthetic hand function for individuals with paralysis.

## Acknowledgements

We would like to thank the research participants for their dedication, hard work, and commitment to the study. We would also like to thank Gracie Hilber and Carleigh May for their help with experiment coordination and data collection. Research reported in this publication was supported by the National Institutes of Neurological Disorders and Stroke of the National Institutes of Health under Award Numbers U01NS123125, UH3NS107714, and T32NS086749. The content is solely the responsibility of the authors and does not necessarily represent the official views of the National Institutes of Health. Data and code related to this study are available on the Data Archive for the BRAIN Initiative (DABI) (https://doi.org/10.18120/65ga-ke41) and GitHub (https://github.com/pitt-rnel/static-dynamic-grasp-force).

---

**Supplementary Figure 1:**
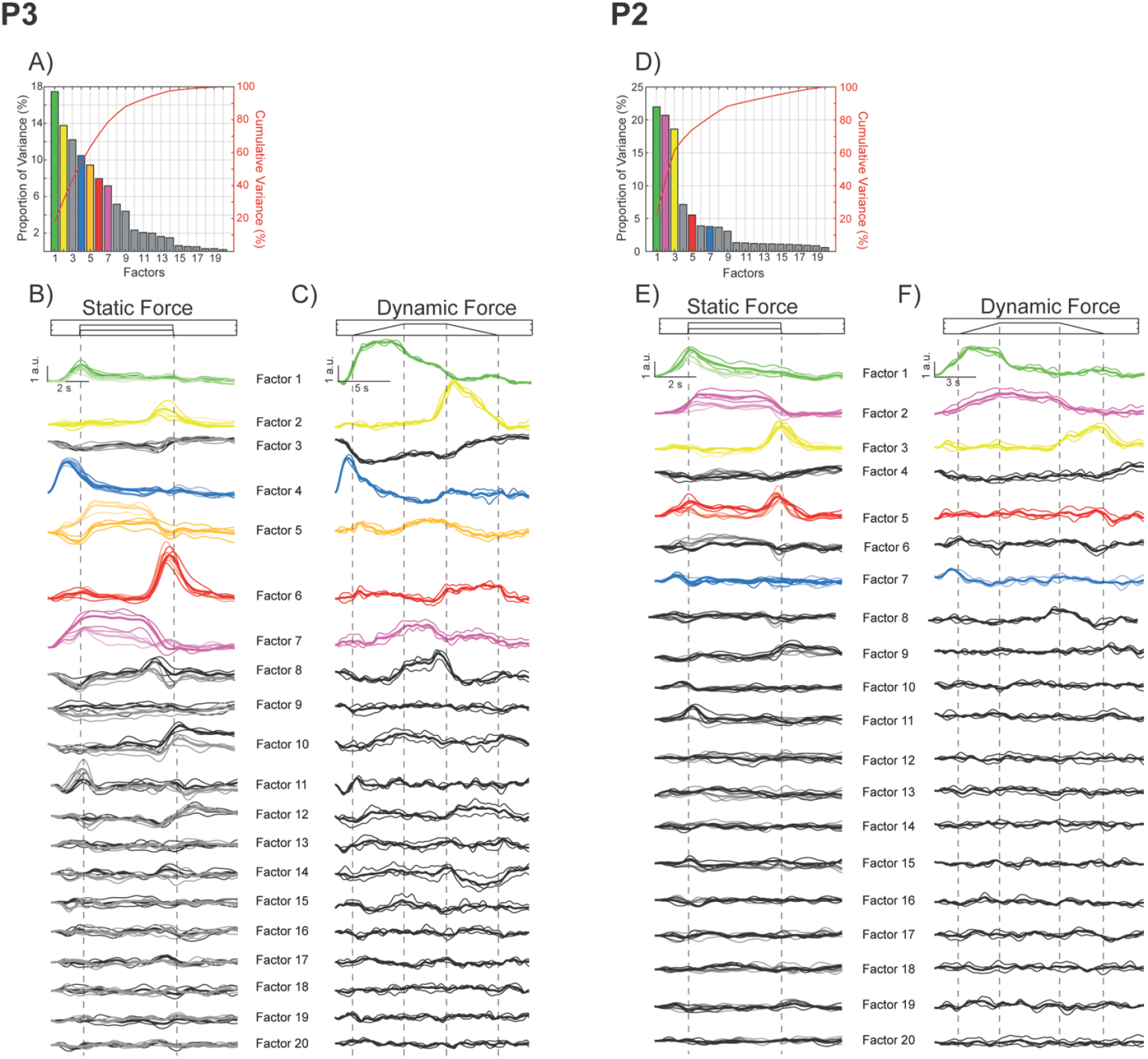
Factor analysis of MC activity: A & D) Skree plots representing variance explained for each factor. B,C,E,F) Projected factors from static force and dynamic force grasping conditions. Each trace represents one session of a GPA aligned dataset. In the static force condition, lighter color traces are from low force grasps and darker traces are from high force grasps. Factor traces with distinguishing color appeared to be modulated during specific phases of the grasp across all conditions.

**Supplementary Figure 2:**
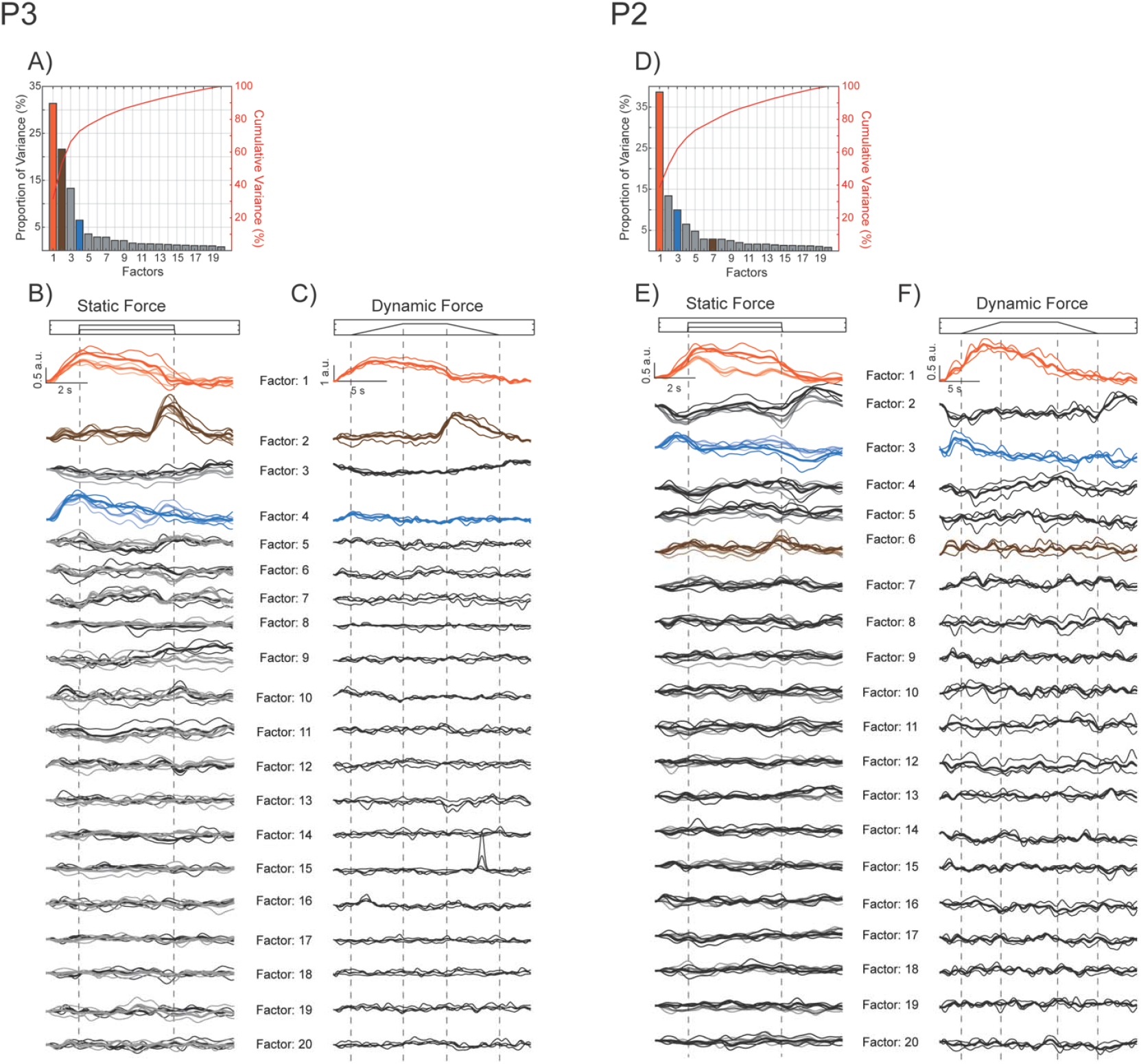
Factor analysis of SC activity: A & D) Skree plots representing variance explained for each factor. B,C,E,F) Projected factors from static force and dynamic force grasping conditions. Each trace represents one session of a GPA aligned dataset. In the static force condition, lighter color traces are from low force grasps and darker traces are from high force grasps. Factor traces with distinguishing color appeared to be modulated during specific phases of the grasp across all conditions.

